# msiFlow: Automated Workflows for Reproducible and Scalable Multimodal Mass Spectrometry Imaging and Immunofluorescence Microscopy Data Processing and Analysis

**DOI:** 10.1101/2024.08.24.609403

**Authors:** Philippa Spangenberg, Sebastian Bessler, Lars Widera, Jenny Bottek, Mathis Richter, Stephanie Thiebes, Devon Siemes, Sascha D. Krauß, Lukasz G. Migas, Siva Swapna Kasarla, Prasad Phapale, Jens Kleesiek, Dagmar Führer, Lars C. Moeller, Heike Heuer, Raf Van de Plas, Matthias Gunzer, Oliver Soehnlein, Jens Soltwisch, Olga Shevchuk, Klaus Dreisewerd, Daniel R. Engel

## Abstract

Multimodal imaging by matrix-assisted laser desorption ionisation mass spectrometry imaging (MALDI MSI) and immunofluorescence microscopy holds great potential for understanding pathological mechanisms by mapping molecular signatures from the tissue microenvironment to specific cell populations. However, existing open-source software solutions for analysis of MALDI MSI data are incomplete, require programming skills and contain laborious manual steps, hindering broadly applicable, reproducible, and high-throughput analysis to generate impactful biological discoveries across interdisciplinary research fields. Here we present msiFlow, an accessible open-source, platform-independent and vendor-neutral software for end-to-end, high-throughput, transparent and reproducible analysis of multimodal imaging data. msiFlow integrates all necessary steps from import and pre-processing of raw MALDI MSI data to visual analysis output, as well as registration, along with state-of-the-art and newly developed algorithms, into automated workflows. Using msiFlow, we unravel the molecular heterogeneity of leukocytes in infected tissues by spatial regulation of ether-linked phospholipids containing arachidonic acid. We anticipate that msiFlow will facilitate the broad applicability of MSI in the emerging field of multimodal imaging to uncover context-dependent cellular regulations in disease states.

## Introduction

The cellular heterogeneity in tissues has a reciprocal and decisive influence on the microenvironment and enables a balance between homeostasis and inflammation. Intra- and intercellular communication is key in both tissue homeostasis and inflammation, and lipids are emerging as critical regulators and key molecules in these processes^1^. Moreover, the lipid landscape was recently defined as a feature of immune cell identity^2^. Upon inflammation, neutrophils, the most abundant circulating white blood cell subset, readily infiltrate into and navigate through the tissue. Herein, they respond to a plethora of signals from the microenvironment by synthesis of lipid messengers, such as arachidonic acid (AA) and the oxidised metabolites prostaglandins and leukotrienes, influencing recruitment, phenotype, and function of neutrophils^3–5^. In urinary tract infection (UTI), the third most common bacterial infection in humans^6,7^ induced by uropathogenic *Escherichia coli* (UPEC)^8^, neutrophils and urothelial cells are critical cell populations which provide the first immunological barrier for the containment of infection^9–11^. Data on lipidomic adaptations of urothelial cells and neutrophils in UTI are missing, as algorithms assigning lipids to specific cell populations in tissues are scarce, hindering novel insights into the decisive role of lipids in regulating mechanisms of inflammation and resolution.

In order to decipher the cellular interplay and the behaviour of specific cell populations in tissues, methods are required enabling in-depth lipidomic profiling with spatial resolution in the micrometer range. Immunofluorescence microscopy (IFM) efficiently determines the distribution of various cell types in tissue niches with high spatial resolution^12^. In contrast matrix-assisted laser desorption ionisation mass spectrometry imaging (MALDI MSI) with laser-induced postionisation (MALDI-2) provides a label-free technology to investigate the spatial distribution of a large number of lipids and metabolites that are predominantly inaccessible by IFM^13^. Moreover, transmission-mode MALDI-2 (t-MALDI-2) with a high-resolution pixel size of 1 µm was introduced recently^14,15^. Thus, combining IFM and MALDI MSI would enable the assignment of the spatial lipidome to specific cell populations. A recent study integrated multiplex IFM and MSI to map myeloid heterogeneity in its metabolic and cellular context^16^. However, analysis of high-dimensional MALDI MSI data and image co-registration to IFM remains challenging due to the lack of algorithms and complete workflows, that allow reproducible and automated pre-processing, analysis and visualisation of MALDI MSI data^17,18^. As technology advances to achieve higher resolutions, data sizes are increasing, thereby further complicating data handling. Therefore, most commercial software solutions offer a user interface to pre-processed data with reduced size. Although this enables interactive data visualisation and analysis, it offers limited transparency and control over data pre-processing and data quality. In contrast to commercial software, which often remains a black-box for users, existing open-source software offers incomplete solutions, as it is mostly designed for specific tasks (e.g. individual pre-processing steps, image registration, analysis or visualisation) and often requires programming skills or contains laborious manual steps (e.g. manually selecting off-/on-tissue regions)^18–24^. As a result, customised data analysis pipelines are constructed from a pool of open-source packages, in-house developed or commercially available software, hindering reproducible and high-throughput analyses and deterring non-expert users.

In this study, we aimed to bridge this gap by developing msiFlow, an open-source software that integrates all steps from import and pre-processing of raw multimodal and multi-vendor imaging data to registration, analysis and visualisation in automated workflows. The workflows operate fully automatically on all major operating systems. By employing msiFlow in a clinically relevant proof-of-concept study, we provide novel insights into the spatial lipidomic interactome in UTI, revealing a hitherto unknown heterogeneity of neutrophils important for the immune response against invading pathogens.

## Results

### The msiFlow software

High-dimensional molecular imaging through MALDI MSI holds great potential for comprehensive spatial mapping of the cellular heterogeneity in tissues and deciphering complex molecular interactions within the tissue microenvironment. However, the lack of open-source and easy-to-use software for automated MSI data processing and analysis greatly complicates reproducible and precise mapping of molecular landscapes *in situ*. To solve this problem, we integrated, optimised and further developed existing bioinformatic methods for data pre-processing, registration, analysis and visualisation into msiFlow, a collection of automated Snakemake workflows enabling reproducible and scalable analyses^25^. We applied msiFlow using a correlative imaging approach consisting of high resolution (t-)MALDI-2 MSI and IFM in an experimental model of UTI. For this purpose, consecutive mouse bladder sections of 8 µm were measured by IFM, t-MALDI-2 MSI and MALDI-2 MSI with a pixel size of 0.2 µm, 2 µm and 5 µm respectively (Fig. 1A). t-MALDI-2 data were measured by orbitrap and MALDI-2 MSI by time-of-flight (TOF). The generated multimodal imaging data were processed and analysed by msiFlow.

**Fig. 1:**
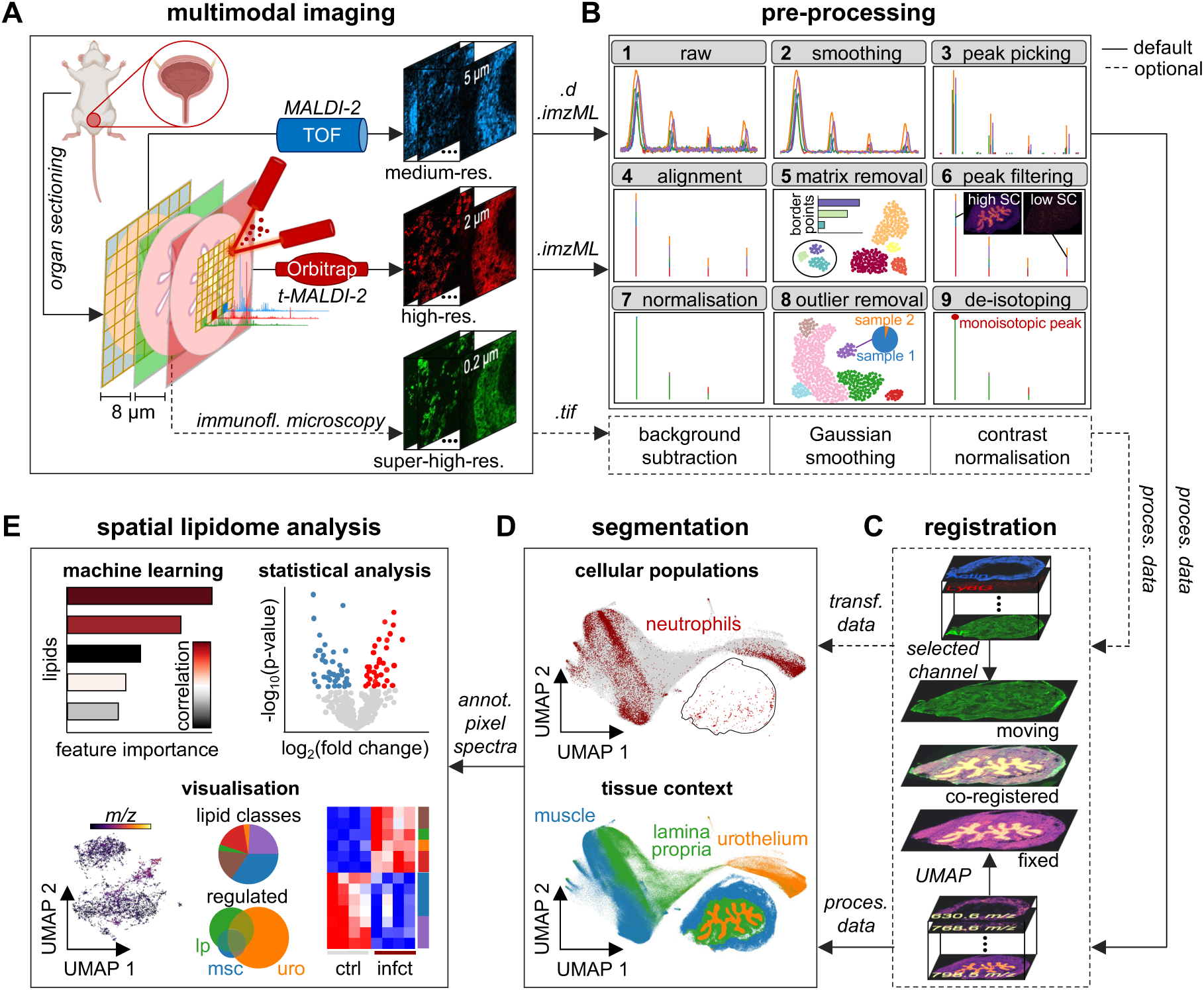
Workflow for correlative multimodal mass spectrometry imaging and immunofluorescence microscopy data. **(A)** Consecutive mouse bladder sections of 8 µm are measured by immunofluorescence microscopy (green), matrix-assisted laser desorption ionisation combined with laser-induced postionisation mass spectrometry imaging (MALDI-2 MSI) (blue) and transmission-mode MALDI-2 MSI (red) with a lateral resolution of 0.2 µm, 5 µm and 2 µm respectively. MALDI-2 MSI data are measured by time-of-flight (TOF) and t-MALDI-2 data by orbitrap. **(B)** The pre-processing workflow takes the raw MSI data in the common data formats as input and outputs the processed data (proces. data). **(B1)** The workflow integrates raw files as input. **(B2)** Spectra are Savitzky-Golay smoothed. **(B3)** Peaks with a user-defined signal-to-noise are selected. **(B4)** A common mass-to-charge vector is formed by a kernel-based clustering approach. Peaks are aligned by nearest-neighbour mapping. **(B5)** UMAP clusters which are connected to most pixels of the border of the measured area are considered matrix/off-tissue clusters. **(B6)** Ions with low spatial coherence (SC), a measure for ion’s informativeness, are removed. **(B7)** Data are normalised to the total ion current (TIC) or median fold change (MFC). **(B8)** UMAP clusters in which most pixels originate from one sample are considered outliers. **(B9)** De-isotoping exploits the theoretical isotope pattern. The software contains an optional step for IFM pre-processing and image registration to combine MSI and IFM. The IFM pre-processing step removes noise from the microscopy images by rolling-ball background subtraction and Gaussian smoothing, followed by contrast normalisation using percentile stretching. This suppresses extremely high/low intensities to enhance image contrast. **(C)** The registration step combines MSI and IFM data. Therefore, a selected channel from IFM (here the autofluorescence image) is used as moving image and the UMAP image from MSI is used as fixed image. The learned transformation is applied to all other IFM image channels which results in transformed data. **(D)** Imaging data are segmented and visualised by UMAP. Each point in the UMAP visualisation represents one MSI data point/pixel. Segmentation of the Ly6G IFM image indicates the distribution of neutrophils (top). The MSI data is segmented into its main tissue regions (bottom). **(E)** Lipidomic signatures of segmented regions and cell populations are extracted. Cell-specific changes between segmented regions are extracted by machine learning-based classification and correlation (top left) and statistical analysis (top right). The distribution of the most important lipids for the classification model are visualised by UMAP (bottom left). The lipidomic changes in the segmented tissue regions are illustrated in Venn diagrams (bottom middle). The regulated lipid classes are depicted in pie charts (bottom middle) and heatmaps (bottom right). Immunofluorescence (immunofl.), processed data (proces. data), transformed data (transf. data), annotated pixel spectra (annot. pixel spectra), lamina propria (lp), muscle (msc), urothelium (uro), control (ctrl), infected (infct).

msiFlow contains 7 Snakemake workflows for pre-processing, registration, segmentation, and analysis/visualisation. We have deliberately divided the software into 7 main workflows to make the application highly flexible and modular. This modular software design enables easy integration of individual workflows of msiFlow into existing analysis pipelines. It is also possible to combine multiple workflows of msiFlow into one workflow via Snakemake. All workflows are integrated into a Docker^26^ image enabling easy-to-use execution on all major operating systems. Each workflow can be run fully automatically through one command in the terminal. Parameters used by msiFlow are defined in one configuration file and can be adjusted by the user depending on the instrument’s setting (e.g. mass and spatial resolution) and preferred methods. msiFlow also provides a browser-based interface to adjust the parameters and run the workflows (Supplementary Fig. 1). A detailed description of all parameters and the configurations used in this study for MSI pre-processing is provided in Supplementary Table 1 and on GitHub for all workflows.

msiFlow includes a MSI pre-processing workflow which imports raw MSI files from different vendors (Bruker and Thermo Fisher), processes all files in parallel, and outputs the processed data in the open standard imzML format along with quality control visualisations (Fig. 1B). The workflow contains steps for spectral smoothing, peak picking, peak alignment, matrix removal, peak filtering, normalisation, outlier removal and de-isotoping to generate an endogenous/tissue-origin mono-isotopic peak list (see method details in Methods). We established two UMAP-based clustering approaches to automatically identify and remove off-tissue/matrix pixels and outliers enhancing data quality for subsequent analysis. To this end MALDI-2 MSI data of each sample were reduced to two dimensions by UMAP^27^ followed by HDBSCAN clustering^28^ (Supplementary Fig. 2A). The cluster connected to most border points of the measured area was considered the off-tissue cluster (Supplementary Fig. 2B). Clusters with high correlation to the off-tissue cluster were combined to an extended off-tissue cluster (Supplementary Fig. 2C) and post-processed to remove isolated objects and fill holes (Supplementary Fig. 2D). With this unsupervised approach we prevent the need of defining known matrix peaks beforehand which are often not consistent throughout all datasets. However, msiFlow also implements a supervised approach based on known matrix peaks. For identification of outliers, data of all samples were reduced to two dimensions by UMAP followed by HDBSCAN clustering (Supplementary Fig. 3A) and sample-specific clusters (SSC) were identified in which 70% of pixels originate from one sample (Supplementary Fig. 3B). Finally, samples in which most pixels were SSC pixels were considered sample outliers (Supplementary Fig. 3C-D).

In addition to MSI pre-processing, msiFlow provides a workflow for IFM pre-processing and image co-registration to combine MSI and IFM data. In the pre-processing workflow, noise is removed from the images by rolling-ball background subtraction and Gaussian smoothing, followed by contrast normalisation (Fig. 1B bottom). Here extremely high/low intensities are suppressed to enhance image contrast. After data pre-processing, MALDI-2 MSI data are combined with the IFM data through image co-registration in which a transformation aligns a moving to a fixed image (Fig. 1C). Several methods have been developed to generate one image out of a MALDI MSI dataset to spatially visualise molecular differences and similarities^29^. Here we used UMAP to reduce the MALDI-2 MSI data to one dimension. Through this approach, we receive one value for each pixel spectrum which can be visualised as a greyscale image. This UMAP image represents the main tissue structure. From IFM we used the autofluorescence (AF) image as it similarly represents the main tissue structure. Registration to the MALDI-2 MSI data was performed using symmetric normalisation implemented in the Advanced Normalisation Tools (ANTs) library^30^. To account for tissue deformations between the consecutive sections, the workflow uses rigid, affine, and deformable transformation with mutual information as the optimisation metric.

From the pre-processed (and registered) data, regions of interest (ROIs) are extracted through two segmentation workflows (one for MSI and one for IFM data) (Fig. 1D). The MSI segmentation workflow includes state-of-the-art dimensionality reduction methods (PCA, t-SNE^31^, UMAP) and clustering algorithms (k-means, spatial k-means^32^, HDBSCAN, hierarchical, gaussian mixture models) to perform unsupervised segmentation which extracts the main tissue context. The IFM segmentation workflow uses a thresholding-based approach to segment specific markers.

For analysing the ROIs msiFlow contains 3 analysis workflows. The first analysis workflow is designed to identify and compare molecular changes in different ROIs (e.g. tissue regions) between two groups by applying statistical analysis. The workflow outputs volcano plots, pie charts, Venn diagrams and heatmaps of the regulated lipids (Fig. 2E right). The second analysis workflow unravels molecular signatures of the ROIs (e.g. cell populations) by using machine learning-based classification (e.g. tree-based classifiers such as AdaBoost, LightGBM^33^, XGBoost^34^), explainable AI methods (e.g. shapely additive values (SHAP))^35,36^ and correlation (e.g. Spearman and Pearson) (Fig. 2E top left). The third analysis workflow applies a UMAP-based clustering approach to reveal molecular heterogeneity in the ROIs (e.g. cell populations) and plots the heterogeneous lipid signals in UMAPs (Fig. 2E bottom left).

**Fig. 2:**
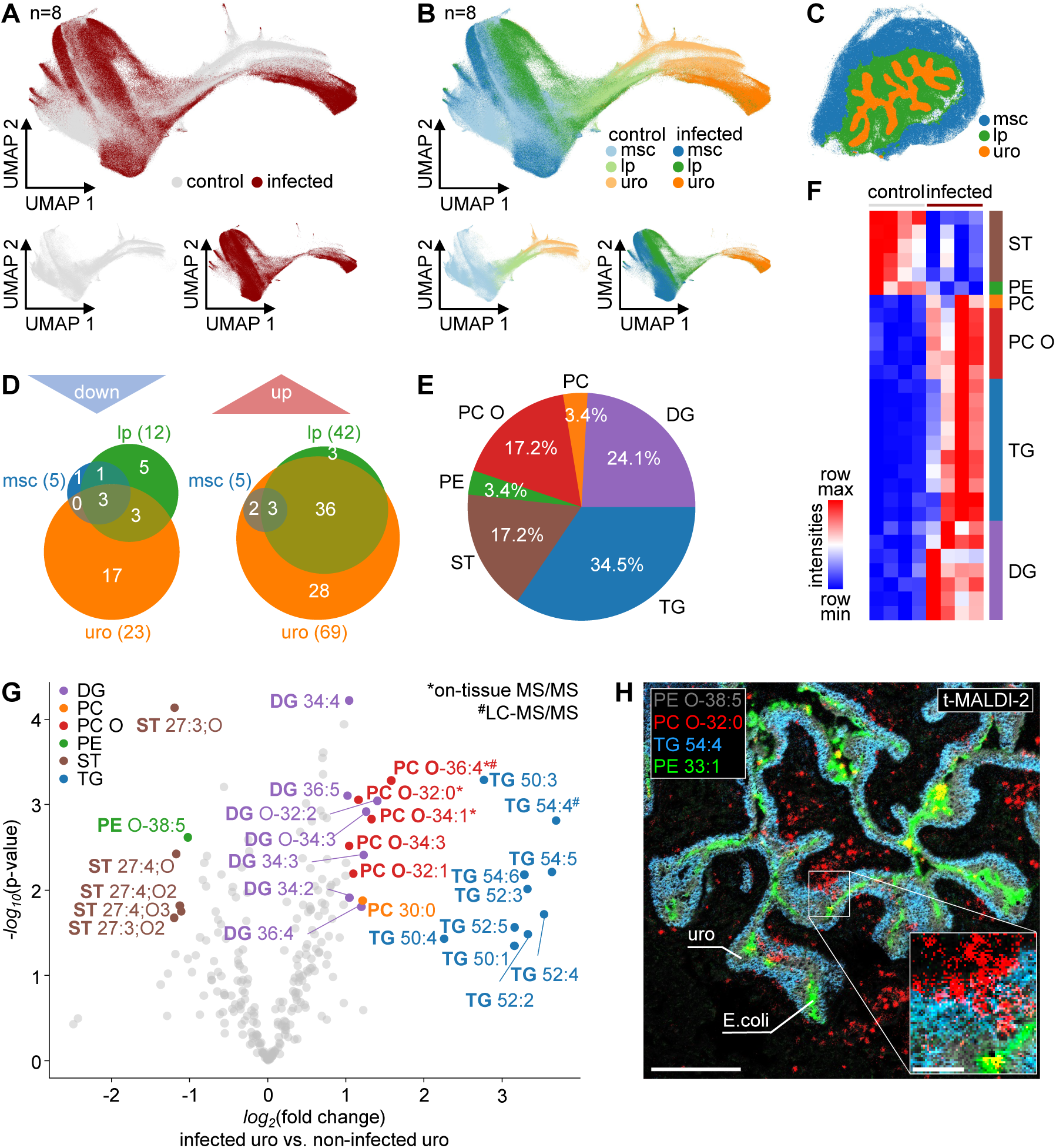
Significant lipidomic adaptations in the urothelium upon infection. **(A)** UMAP embedding of MALDI MSI data of 4 control (grey) and 4 infected (red) bladder sections (n=8). **(B)** UMAP embedding colour-coded according to the muscle (msc), lamina propria (lp) and urothelium (uro) in the control (light-coloured) and infected (dark-coloured) tissue sections. **(C)** Spatial mapping of the UMAP clusters shown in B in an infected tissue section. **(D)** Average spectra of the different tissue regions (msc, lp and uro) were extracted and statistically analysed. Venn diagrams revealing the number of significantly downregulated (left) and upregulated (right) lipids (p-value < 0.05 and |*log_2_*(FC)| > 0.5). **(E)** Pie chart showing the percentage of lipid classes regulated in the infected urothelium. **(F)** Heatmap showing the average intensity of regulated lipids (p-value < 0.05 and |*log_2_*(FC)| > 1) in the urothelium for each sample. **(G)** The significantly regulated lipids (p-value < 0.05 and |*log_2_*(FC)| > 1) in the urothelium are labelled in the Volcano plot. Lipids validated by on-tissue MS/MS are marked with *, and by LC-MS/MS are marked with ^#^. Statistical significance was tested by Welch’s t-test for normally distributed populations and Wilcoxon rank-sum test for non-normally distributed populations. n=8 (4 control versus 4 infected). **(H)** Spatial distribution of regulated lipids and *E.coli* (PE 33:1) in an infected bladder measured by t-MALDI-2 MSI. Scale bars indicate 300 µm and 50 µm in the zoom. Diglyceride (DG), phosphatidylcholine (PC), ether-linked phosphatidylcholine (PC O), phosphatidylethanolamine (PE), sterole (ST), triglyceride (TG).

### Spatial lipidomic changes in the infected urinary bladder

We applied msiFlow to MALDI MSI data of the clinically relevant model of UTI to reveal niche-specific molecular changes upon infections with UPEC. To segment the urinary bladder into the main tissue compartments, we applied dimensionality reduction by UMAP followed by HDBSCAN clustering with manual refinement to the MALDI MSI data. To indicate the molecular changes in the different tissue regions, MS spectra from control and infected samples were visualised and compared by the 2D UMAP embedding (Fig. 2A). Annotation of the spectra to specific tissue areas, i.e., lamina propria (lp), muscle (msc) and urothelium (uro), indicates specific lipidomic signatures in the different tissue regions in control and infected samples (Fig. 2B). In addition, the UMAP visualisation indicates specific tissue layers, such as the urothelium, with strong changes upon infection. Spatial mapping of the UMAP representation visualises the different tissue regions (Fig. 2C). Next, we generated tentative lipid annotations for the *m/z* values by using the bulk structure search from the LipidMaps Website (www.lipidmaps.org) and searched for expected lipid classes and [M+H]^+^, [M+Na]^+^ and [M+K]^+^ precursor ions with a mass tolerance of +/- 0.01 *m/z*. For further analysis, *m/z* signals without potential lipid matches were filtered out in order to eliminate non-lipid peaks (e.g. in-source fragments or chemical background). Then we compared the lipidomic changes separately in the urothelium, lamina propria and muscle. Therefore, we performed statistical analysis of the mean intensities of urothelial, lamina propria and muscle pixels of infected vs. control samples and performed MALDI MSI in data-dependent acquisition (DDA) mode to validate the lipids (Supplementary Table 2). This analysis indicated the strongest lipidomic alterations in the urothelium, the site of bacterial tissue entry, across various lipid classes (Fig. 2D-E). The main altered lipid classes included triacylglyerols (TG), diacylglycerols (DG) and phosphatidycholines (PC), all of which were upregulated in the infected urothelium (Fig. 2F-G). Among other lipids, TG 54:4 was exclusively expressed in the urothelium (Fig. 2H). In contrast, PC O-32:0 was not only expressed in the urothelium, but also in the lamina propria close to the urothelial expression of TG 54:4. High resolution MSI (2 µm) by t-MALDI-2 revealed the distribution of the UPEC infection by PE 33:1 (704.52 *m/z*), an odd chain fatty acyl PE known to be highly expressed in *E.coli*^37^ (Fig. 2H). The t-MALDI-2 measurement further indicated the presence of ramified cells (PC O-32:0) in the bladder tissue and phagocytosis of bacteria in TG 54:4-rich urothelial areas by those highly ramified cells.

### Identification of lipidomic signatures of neutrophils

Among others, neutrophils are critical immune cells during bacterial infections and efficiently migrate into infected tissue areas to phagocytose bacteria^10^. We detected Ly6G^+^ neutrophils in the lamina propria around blood vessels and in the infected urothelium suggesting directed migration towards the infection (Supplementary Fig. 4). Microscopy also indicated downregulation of CXCR2 in the urothelium, suggesting niche-specific desensitisation of this chemokine receptor at the site of infection (Supplementary Fig. 4).

To unravel the niche-specific lipidomic signature of neutrophils, we used our multimodal imaging workflow and performed IFM and MSI on the timsTOF fleX instrument on consecutive tissue sections one day after infection. Ly6G, a specific marker for neutrophils, was used to localise neutrophils across the tissue and actin, expressed in the muscle tissue, was used to demarcate the organ boundaries to the outside tissue areas and the lamina propria (Fig. 3A). For image co-registration, the AF channel from IFM was used as moving and the UMAP representation from MSI as fixed image (Fig. 3B). The learned transformation was applied to all IFM image channels. The registration result was validated by the Jaccard index for the overlap between transformed urothelial mask from IFM and MSI (average 0.8 Jaccard). Registered Ly6G images were segmented to annotate the pixel spectra according to neutrophil-rich areas. Then a binary tree-based classifier was trained with the Ly6G-annotated pixel spectra to extract lipidomic signatures of neutrophils (Fig. 3C). The most important lipids to classify Ly6G^+/-^ pixel spectra were several ether-linked PCs (Fig. 3D). For the classification model, lipids that are anti-correlated with neutrophils or present in very low abundance in neutrophils are as important as lipids which are highly correlated to neutrophils or present in very high abundance. To distinguish between lipids that are highly and lowly abundant in neutrophils, the bar plot is colour-coded according to Pearson’s correlation (Fig. 3D). The lipid with the highest feature importance and correlation for neutrophils was PC O-36:4 (Fig. 3E). LC-MS/MS (Supplementary Fig. 5) and *in situ* MS/MS (Supplementary Fig. 6) indicated that PC O-36:4 contains the esterified AA.

**Fig. 3:**
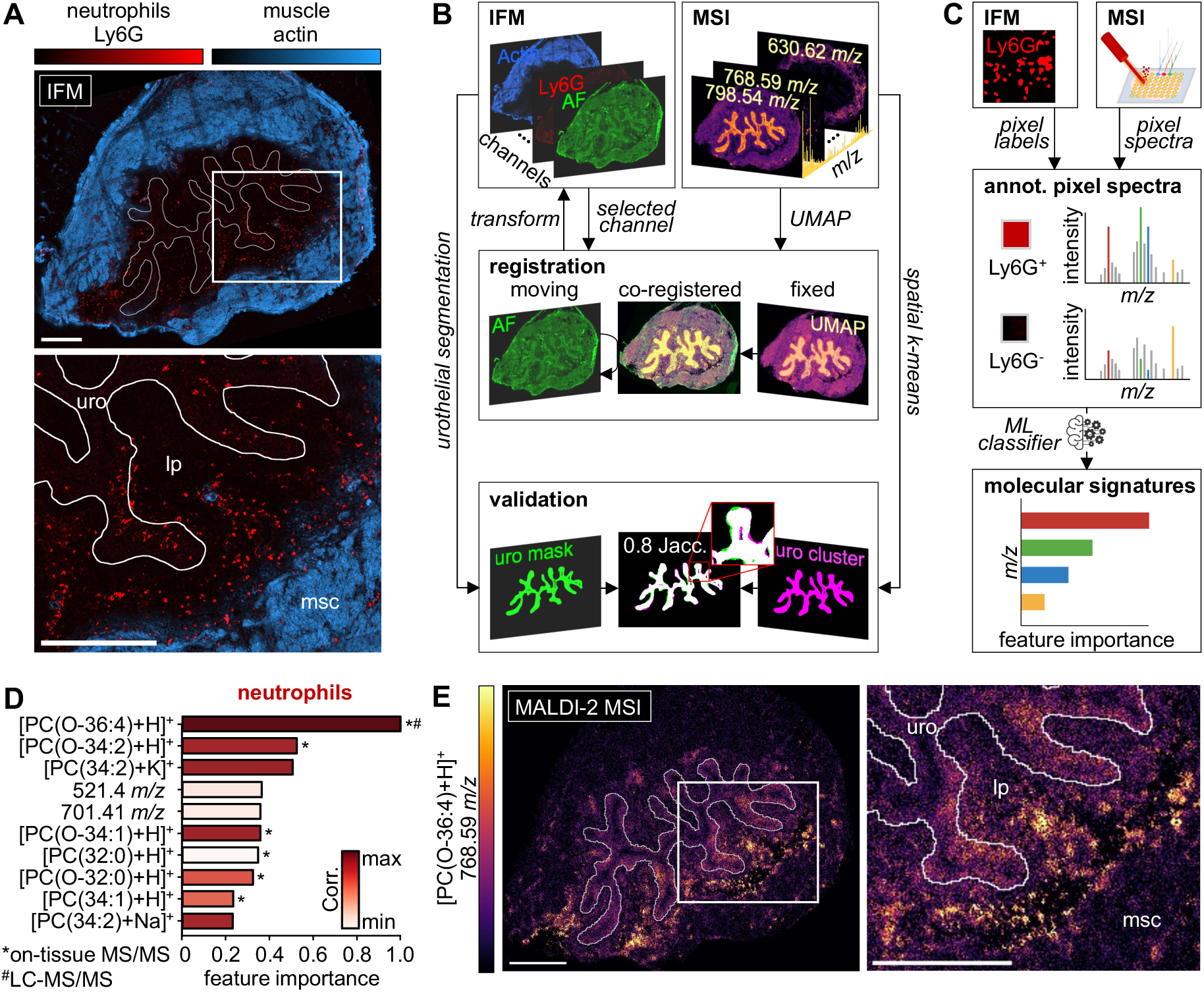
Image co-registration and machine learning-based classification extract spatial lipidomic signatures of neutrophils. **(A)** IFM of Ly6G for neutrophils (red) and actin for smooth muscle cells (blue) was performed on consecutive bladder sections to the MALDI MSI sections. **(B)** IFM images were registered to the MALDI MSI images via symmetric normalisation implemented in the Advanced Normalisation Tools (ANTs) library. The AF image from IFM was used as moving image and the UMAP image from MSI served as fixed image. The learned transformation was applied to all IFM image channels. We validated the registration result by the Jaccard index for the overlap between transformed urothelial mask from IFM (green) and MSI (magenta). We applied spatial k-means clustering on the MSI data and selected the cluster containing the urothelium. The urothelial mask from microscopy was manually created. **(C)** Segmentation of the registered Ly6G images was performed to annotate the pixel spectra into neutrophil/non-neutrophil pixels. Then a machine learning-based classification model was trained with the annotated pixel spectra for neutrophils. **(D)** Feature importance ranking revealing the top 10 lipids which are most important for the classifier to distinguish neutrophil and non-neutrophil pixels. The bars are colour-coded according to Pearson’s correlation coefficient. While a high correlation indicates high co-localisation, a low correlation indicates no co-localisation. We identified several ether-linked phospholipids, which were further characterised by on-tissue MS/MS (marked with *), to be highly correlated to neutrophils distribution. The top lipid was also validated by LC-MS/MS (marked with ^#^). **(E)** Ion image of the top lipid identified for neutrophils distribution on the consecutive section to the IFM section shown in A. Immunofluorescence microscopy (IFM), mass spectrometry imaging (MSI), autofluorescence (AF), Jacc. (Jaccard), annot. (annotated), machine learning (ML), Pearson’s correlation (corr.), urothelium (uro), lamina propria (lp), muscle (msc). Scale bars indicate 500 µm.

To validate our results, we performed MALDI-2 MSI from bone marrow (BM)-derived neutrophils and applied our data pre-processing workflow (Supplementary Fig. 7A). To extract the main peaks from BM neutrophils, we filtered the peaks based on their spatial coherence and average intensity. The filtered peaks include the top 4 lipids (M+H) for bladder neutrophils. These lipids are expressed at a very high level in BM neutrophils, indicating that our imaging workflow identifies cell population-specific lipids (Supplementary Fig. 7B).

### Lipidomic heterogeneity of neutrophils in the infected urinary bladder

Neutrophils can adopt specific maturation states in secondary lymphoid organs such as the bone marrow and spleen, thus establishing significant heterogeneity that is important for defence against pathogens. However, little is known about this heterogeneity in infected organs, although certain cellular states may represent specific adaptations to the tissue environment and infectious conditions. To reveal the lipidomic heterogeneity of neutrophils in the infected urinary bladder, we used msiFlow. Here registered Ly6G images were first segmented and annotated according to the tissue region (urothelium, lamina propria and muscle) in which they are localised (Fig. 4A-B), followed by dimensionality reduction by UMAP (Fig. 4C). Next HDBSCAN clustering of MSI spectra of Ly6G^+^ pixels was performed, generating 3 clusters (Fig. 4D-E). A classification model was trained on the clustered Ly6G^+^ pixel data to identify the lipids with the strongest feature importance and SHAP values. We found that PC O-36:4 was the most important lipid for the classification model based on the feature importance (Fig. 4F). Mapping of the clusters to the different tissue compartments revealed that neutrophils in cluster 0 are mainly located in the urothelium by 83%, neutrophils in cluster 1 are mainly located in the lamina propria by 88% and neutrophils in cluster 2 are equally distributed in the lamina propria (52.7%) and muscle (46.4%) (Fig. 4G). In addition, binary classifiers were trained for each class which revealed class-specific lipidomic signatures (Fig. 5H-J). The distribution of the top lipids showed strong expression of DG 34:2 (575.5 *m/z*) in cluster 0 (Fig. 5H), PC O-36:4 (768.59 *m/z*) in cluster 1 (Figure 4J) and SM 42:2;O2 (851.54 *m/z*) in cluster 2 (Fig. 4I). These data demonstrate the spatial expression of PC O-36:4, which contains the immunologically important metabolite AA, in neutrophils in different tissue niches in the urinary bladder. msiFlow resolved the lipidomic heterogeneity of neutrophils in infected tissue areas, shedding new light on the tissue-specific lipidomic adaptations of neutrophils and providing new possibilities for the adaptations of an immunological response in tissues.

**Fig. 4:**
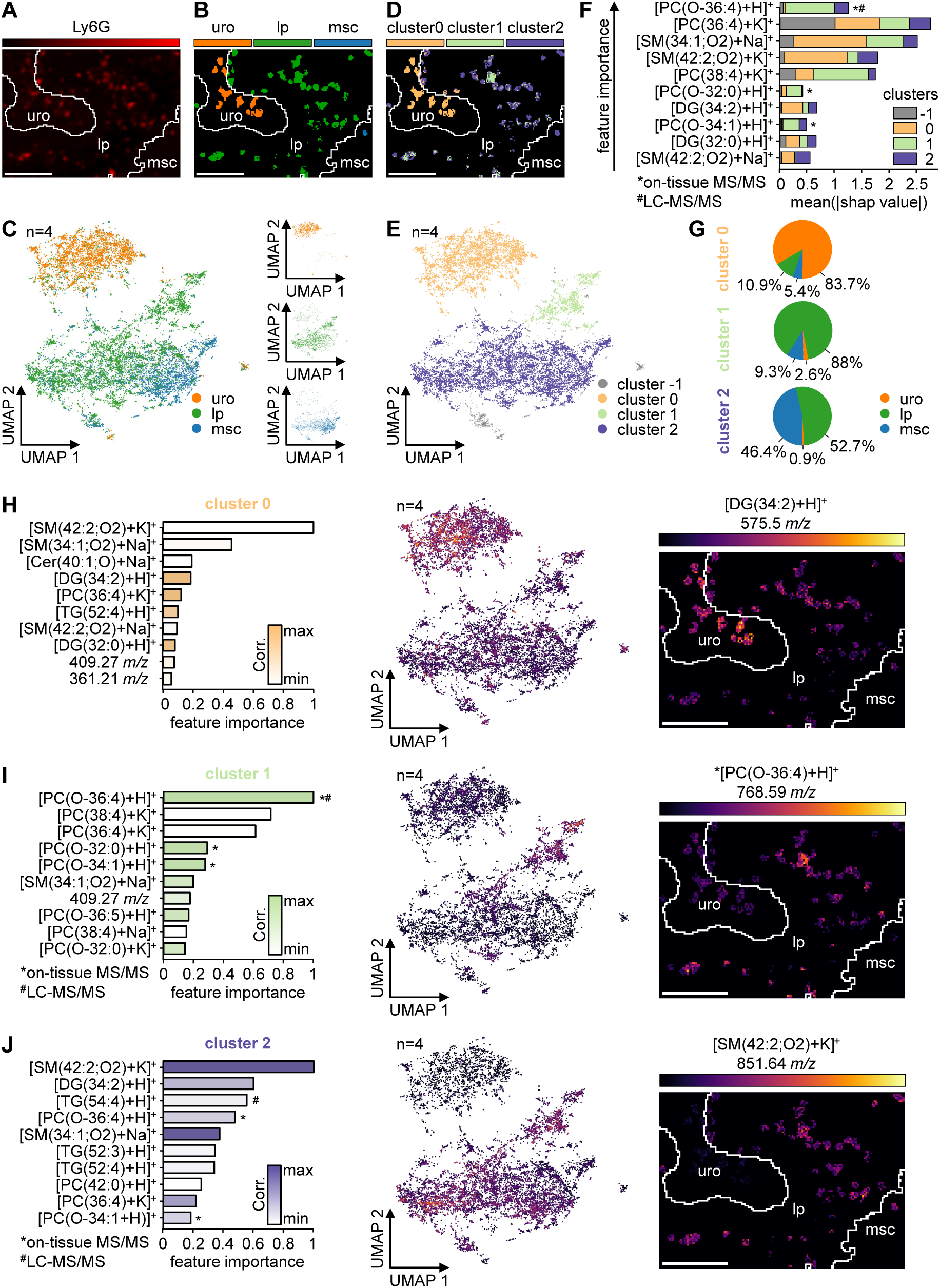
Lipidomic heterogeneity of neutrophils. **(A)** Registered Ly6G image showing the distribution of neutrophils in the uro, lp and msc. **(B)** Segmented Ly6G image, colour-coded according to the different tissue regions. **(C)** UMAP of Ly6G^+^ pixel spectra in 4 infected bladder samples (n=4) colour-coded according to the different tissue regions. **(D)** HDBSCAN clustering was performed on the UMAP embedding of Ly6G^+^ pixel spectra. The spatial distribution of the 3 main clusters is shown in the image of an infected bladder region. **(E)** HDBSCAN clusters shown in the UMAP embedding of Ly6G^+^ pixels. Cluster −1 indicates noise pixels. **(F)** A XGBoost multi-class classifier was trained on the annotated clustered pixel spectra to extract the mean SHAP values of the 10 most important lipids based on the feature importance. The top 10 lipids are shown in ascending order according to their feature importance. **(G)** Distribution of cluster pixels in the different tissue regions. **(H-J)** A XGBoost binary classifier was trained for each cluster to reveal cluster-specific lipidomic signatures. The resulting top 10 lipids based on their feature importance are shown in the bar plots (left) and coloured according to their Pearson’s correlation. Signal intensities of lipids with the highest feature importance and positive Pearson’s correlation for cluster 0 **(H)**, cluster 1 **(I)** and cluster 2 **(J)** shown in the UMAP (middle) and a spatial heat map (right). Urothelium (uro), lamina propria (lp) and muscle (msc). Scale bars indicate 200 µm. n=4 (UPEC infected).

## Discussion

In this study msiFlow, a powerful open-source end-to-end software for automated pre-processing, analysis, visualisation, and registration of MALDI MSI and IFM data was generated. Using msiFlow, we clarified the cell-specific lipidomic adaptations of urothelial cells and neutrophils in UTI at a high spatial resolution and identified specific ether-linked PCs and the AA metabolism by neutrophils in certain tissue niches.

With msiFlow, we address the current lack of complete, automated, and open-source MSI software by integrating all necessary steps, from raw data import to multivariate analysis and visual output, as well as registration into msiFlow. All steps are automated in Snakemake workflows, which enables parallel data processing for high-throughput analyses. All workflows can be run via a single command in the terminal on all major operating systems and do not require complicated package installations, as we incorporated all workflows into a Docker image, making it broadly applicable and suitable for non-programmers compared to common software which only runs on Windows (e.g. SCiLS Lab, LipoStar and msiQuant^38^) or requires programming skills (e.g. Cardinal). Nevertheless, msiFlow provides full flexibility to define all desired steps and preferred methods to be executed by specifying a manageable number of parameters in one configuration file. In addition, msiFlow is fully developed in the open-source programming language Python, as opposed to the widely used proprietary programming language MATLAB. Thus, msiFlow is more affordable and accessible than commercial or proprietary software developed in MATLAB^16,22,39^. Unlike most software that only accepts data in imzML format^21,22,38–41^, msiFlow accepts raw timsTOF data in addition to imzML, preventing the need for additional software for data parsing and the dependence on manufacturer-specific software.

We have made special efforts to develop a complete pre-processing pipeline to filter the data into an endogenous/tissue-origin mono-isotopic peak list for downstream analysis. Simple peak picking is often not sufficient to filter out non-informative or exogenous signals (e.g. from matrix, isotopes and chemical noise) from raw spectra^42^. Such unfiltered data can lead to prediction models learning from non-biological meaningful signals. Therefore, our workflow includes a peak filtering step in addition to standard peak picking. Our workflow performs pixel-wise peak-picking and utilises the spatial information for peak filtering by computing the spatial coherence metric^43^, presenting a significant improvement over common MS software that uses single mass spectra without considering spatial information^44^. In addition, we have developed two UMAP-based clustering approaches that sufficiently determine matrix/off-tissue pixels and outliers in datasets fully automatically, which was previously often done manually (e.g. manual selection of specific regions in SCiLS Lab). The automated matrix detection approach is applicable for whole-sample measurements due to its reliance on tissue architecture and borders and the outlier detection approach is effective for multi-sample datasets as it relies on spectral similarity across multiple samples. Our workflow also provides visual results after each pre-processing step, offering transparency and the possibility of quality control compared to commercial software.

msiFlow not only enables end-to-end analysis of MSI data, but also registration with IFM for multimodal imaging, complementing existing open-source software which either does not include registration^41^ or focuses on registration without data pre-processing and analysis^18^. In this study we registered IFM to high-resolution MALDI MSI to account for the emerging interest in multimodal analyses. Typically, immunohistochemistry of tissue sections is registered to molecular data from MSI and annotated by pathologists^45,46^. However, registration with IFM enables the precise localisation of specific cell populations to characterise the phenotypical and molecular heterogeneity of cells in the tissue context. We deliberately used consecutive sections as the washing steps of the staining procedure of cell-specific molecules strongly reduces the signal-to-noise of the spatial lipidome. Thus, our approach avoids this problem by registering imaging data from consecutive tissue sections. However, we acknowledge limitations in achieving complete cell-specificity due to the usage of consecutive tissue sections and lateral diffusion of lipids from surrounding cells during MSI sample preparation. While our automated registration approach might be less accurate than the popular landmark-based registration selecting features present in both modalities^16,47^ or laser ablation marks from MSI^18^, it does not require manual steps and is not dependent on a post-acquisition pattern. Furthermore, our MSI data of BM-derived neutrophils revealed a substantial abundance of the identified lipids, validating the adequacy of our image registration, segmentation, and feature extraction approach for identifying molecular signals associated with particular cells within tissue sections to a certain degree.

So far, msiFlow does not include an annotation or interpretation step, as we see the main aim of the software in identifying interesting molecular patterns and preliminary candidates, which can be validated by MS/MS in a next step. However, as msiFlow provides a mono-isotopic peak list and contains a mass alignment procedure, tentative lipid annotations for interesting candidates can be made through mass matching to databases, such as LipidMaps, as presented in this study. Due to the modular software design, an annotation routine as well as available software for biological interpretation^48^ can easily be added in the future. Moreover, we anticipate that msiFlow can be scaled to other MSI modalities, such as metabolomics and proteomics.

To demonstrate the strength of our multimodal imaging approach and msiFlow to biological research, we applied msiFlow to lipidomic MSI and IFM data of UPEC-infected bladders. This revealed high abundance of ether-linked PCs in neutrophils, which is in line with previous studies^2,49,50^. Polyunsaturated fatty acids (PUFAs) containing lipids, such as PC O-36:4, strongly influence cellular function via effects on membrane properties, and by acting as a precursor pool for lipid mediators, such as AA. Among others, these mediators are enzymatically converted by cyclooxygenase and lipoxygenase to generate eicosanoids, such as prostaglandins and leukotrienes^4^. These metabolites are critical for neutrophil migration and swarming in the 3-dimensional tissue context^51^. Recently, changes in the composition of PCs were observed during neutrophil differentiation stages, indicating reduced abundance of this PUFA in mature stages of neutrophils^2^. Similarly, we found that PC O-36:4 was most abundant in neutrophils after extravasation in the lamina propria, but not detectable at the site of infection in the urothelium, suggesting metabolic oxidation of AA into leukotrienes to facilitate neutrophil swarming towards the local infection in the urothelium. Besides AA oxidation, the reduced abundance of PC O-36:4 in the urothelium could also indicate neutrophil apoptosis, as it is assumed that ether-linked PCs are important membrane components required for neutrophil survival^52^. Taken together, msiFlow revealed a heterogeneous distribution of PC O-36:4 in neutrophils, indicating a critical and spatial role of PC O-36:4 in neutrophil state and function.

With the advent of our established software and the high-resolution t-MALDI-2 MSI technologies, we found an increased expression of molecules in the population of urothelial cells important for mucus production and modulation of the bacterial colonisation, such as PCs, DGs and TGs^53,54^. Our t-MALDI-2 MSI data revealed an accumulation of TGs in the urothelium close to the bacteria. Interestingly, TGs were also among the lipids with the highest feature importance and correlation to neutrophils in the urothelium. TGs are stored in lipid droplets (LDs), which have been described as sites of eicosanoid synthesis^55^. It is also known that defence molecules, such as Cathelicidin (CAMP) and histones, accumulate on the LDs of challenged cells which makes them more resistant to bacterial species including *E.coli*^56^, potentially through the mechanism of neutrophil extracellular traps (NETs)^56^. Hence, our findings suggest an important role of LDs in supporting the immune defence against UPECs in UTI through eicosanoid synthesis and NET formation.

In conclusion, we established an open-source, platform-independent and vendor-neutral software for automated, end-to-end, transparent, reproducible, and scalable multimodal MSI and IFM image analysis with a low barrier to entry. Using msiFlow, we not only identified lipidomic signatures of neutrophils, validated by MSI of BM neutrophils and in line with previous studies, but also unravelled a hitherto unknown lipidomic heterogeneity of neutrophils in UTI. This is an essential step towards uncovering the context-dependent regulation of leukocytes in inflammatory conditions. The easy usability and completeness of our software will facilitate the applicability of MSI in the emerging field of multimodal imaging.

## Methods

### Animal studies

Female C57BL/6 mice were used throughout the experiments. Animals were purchased from Charles River Laboratories and maintained under specific-pathogen-free conditions in the central animal facility at the University Hospital Essen. The local review board (Bezirksregierung Köln, Landesamt für Natur, Umwelt und Verbraucherschutz NRW in Recklinghausen, Germany) approved the animal experiments.

### Urinary tract infection model

UPEC strain 536 (O6:K15:H31) was cultured for 3 h at 37 °C in LB medium. The bacteria were harvested by centrifugation at 1500 × g for 20 minutes and then the OD(600) was measured. The bacteria were resuspended at a concentration of 10^10^ bacteria/ml in sterile PBS. A mixture of ketamine and xylazine 80/10mg/kg body weight in 150μl PBS was injected intraperitoneally to anesthetise the mice. The infection of the animals was induced by transurethral inoculation of 5 × 10^8^ UPEC in 0.05 ml PBS using a soft polyethylene catheter.

### Immunofluorescence microscopy (IFM)

Urinary bladders were fixed overnight in PLP buffer [pH 7.4, 0.05 M phosphate buffer containing 0.1 M L-lysine, 2 mg/ml sodium periodate, and paraformaldehyde with a final w/v concentration of 1%], equilibrated in 30% sucrose for 24h and stored at −80°C. Bladder tissue was cut into 8 µm thick sections at −20°C using a cryostat. Unspecific binding was blocked by incubation of the sections with PBS containing 1% BSA and 0.05% Triton X-100 for 1 hour. Bladder sections were mounted into MACSwell sample carriers, blocked using a blocking buffer containing 10% BSA and 2% goat serum for 1h at RT before nuclei were counterstained using DAPI-staining solution (Miltenyi Biotec) according to the manufacturer’s recommendations before being placed into a MACSima imaging system. Sections were then incubated with directly conjugated antibodies against Ly6G (1A8, Miltenyi Biotec, PE,1:50), EpCAM (REA977, Miltenyi Biotec, APC,1:50), SMA (REAL650, Miltenyi Biotec, FITC, 1:300) and CXCR2 (SA044G4, BioLegend, PE, 1:50). Acquired pictures were stitched using the preprocessing pipeline in MACS iQ View Analysis Software (Miltenyi Biotec) for downstream analysis.

### Matrix-assisted laser desorption mass spectrometry imaging (MALDI MSI)

Unfixed tissue sections of infected and uninfected bladders were thawed under a gentle stream of dry N_2_ gas. For t-MALDI measurements, the tissue sections were washed with 250 µL of a 150 mM ammonium acetate solution to remove alkali metal salts and dried again under a gentle stream of dry N_2_ gas. 2,5-DHAP matrix was applied onto the tissue section by sublimation in a home-build sublimation device described earlier by Bien et al. ^57^ 1.5 mL of a 20 mg/mL solution of 2,5-DHAP in acetone was filed into the matrix reservoir and heated to about 120 °C which caused the acetone to evaporate. The sample was mounted to the cold side of the sublimation device and kept at around 4 °C at an approximate vertical distance of 6 cm above the matrix reservoir. The sublimation device was evacuated to a pressure of about 5 x 10^-3^ mbar and the sublimation was conducted for 10 min. The samples were measured immediately after sublimation.

All MALDI MSI measurements were carried out in positive-ion mode. MALDI-2-MSI data with a pixel size of 5 µm was acquired with a timsTOF fleX MALDI-2 (QTOF) instrument with microGRID extension (Bruker Daltonics, Bremen, Germany). The MALDI and postionisation laser were operated with a pulse repetition rate of 1 kHz and a delay of 10 µs. The ablation laser power was set to 80% with 25 shots per pixel. The ion detection range was set to *m/z* 300-1500. The TIMS separated MS/MS measurement was acquired with a pixel size of 50 µm with 250 laser shots, a 1/k_0_ range from 1.4-1.8 with N2 as collision gas, a ramp time of 250 ms, an isolation window of 1 Da and 30 eV collision energy.

The t-MALDI-2 MSI data with a pixel size of 2 µm was acquired with a setup which has been described in detail previously ^14,58^. For this a Q Exactive Plus Orbitrap mass spectrometer (Thermo Fisher Scientific, Bremen, Germany) equipped with a modified dual-ion funnel source (Spectroglyph, Kennewick, WA, USA), which enabled transmission mode illumination and MALDI-2 postionisation, was used. Both lasers were operated with a pulse repetition rate of 100 Hz and a delay of 10 µs. A mass resolution of 70000 (defined for *m/z* 200) with a fixed injection time of 250 ms and an ion detection range of *m/z* 350-1500 were used. MALDI-DDA-MSI was used to confirm some of the tentatively annotated lipids with on-tissue MS/MS. For this a similar approach to the one used by Ellis et al. ^59^ was chosen, which alternates full-scan and MS/MS pixel. For this front side illumination was used with a step size in x- and y-direction of 10 µm and 20 µm respectively. For the full scan pixels an ion detection range of *m/z* 550-1500 was used. For DDA MS/MS measurements an isolation window of 1 Da and a fixed first mass of *m/z* 100 with a NCE of 25 were used. The exclusion time was set to 30 s. The mass resolution for both full scan and MS/MS was set to 70000 and the injection time was fixed at 250 ms. The MALDI-DDA-MSI data was analysed using Lipostar MSI (vs. 1.3, Molecular Horizon, Bettona, Italy) and annotated using the Lipid Maps Structure Database (https://www.lipidmaps.org/databases/lmsd).

### Isolation of bone marrow-derived neutrophils

Bone marrow derived neutrophils were obtained using a mouse Neutrophil Isolation Kit (Miltenyi Biotec, 130-097-658) following the manufacturer’s instructions. In brief, the bone marrow of a femur was flushed, and erythrocytes lysed using RBC Lysis Buffer (BioLegend, 420302; 1 min on ice). Cells were then sequentially incubated with Neutrophil Biotin-Antibody Cocktail and Anti-Biotin Microbeads before neutrophils were isolated using a negative selection on a magentic column. The isolated neutrophils were then centrifuged onto slides using a cytospin centrifuge, fixed for 5 min using 4% formaldehyde, washed two times with 500 µL PBS followed by two washes with 500 µL of a 150 mM ammonium acetate solution to remove the PBS. For MALDI-MSI matrix application was carried out as described above and the cells were measured with 50 µm pixel size on the timsTOF fleX MALDI-2 (QTOF).

### Liquid chromatography-tandem mass spectrometry (LC-MS/MS)

Lipids were extracted using methyl tert-butyl ether (MTBE) extraction and analysed with liquid chromatography-tandem mass spectrometry (LC-MS/MS) as reported earlier^60^. In brief, snap-frozen urinary bladder tissues were homogenised in ice-cold IPA:H2O (1:1 v/v; 150μL), spiked with 10μL of SPLASH lipid standard mix, and then subjected to liquid-liquid extraction using MeOH, MTBE, and water solvent mixture with a final ratio of 1:3:1 (v/v/v). The upper organic phase was collected, dried out under a nitrogen gas stream, and reconstituted in 100μL of ACN:IPA:H2O buffer (65:30:5 v/v/v). Resuspended lipid extracts (10μL) were loaded on a reversed-phase ACQUITY UPLC HSS T3 (1.8 μm, 100 × 2.1 mm, Waters Corporation) column and separated using a Vanquish Duo UHPLC-system (Thermo Fisher Scientific, Waltham, MA, USA) with a flow rate of 250 μL/min. The mobile phases consisted of eluent A (H2O:ACN, 40:60 v/v) and eluent B (IPA:ACN, 90:10 v/v) both with 10mM ammonium formate and 0.1% formic acid. All the datasets were acquired independently in negative- and positive-ion mode in a data-dependent manner using Orbitrap Fusion Lumos Tribrid Mass Spectrometer (Thermo Fisher Scientific, Waltham, MA, USA) equipped with a heated electrospray ionisation source. Lipids were annotated and validated with lipid class and molecular-species specific diagnostic fragment ions^61^.

### MSI data pre-processing

MALDI MSI data were pre-processed by custom-designed Python scripts which were automated in a Snakemake^62^ pipeline. The pipeline takes raw timsTOF files as input, processes all files in parallel, and outputs the processed data in imzML format along with various quality control visualisations for each step. All parameters used within this study are listed in Supplementary Table 1. In the first step Savitzky-Golay smoothing^63^ was used to reduce spectral random noise. Centroid spectra were extracted by using the find_peaks function from the SciPy signal processing library with default parameters. Peaks with a signal-to-noise ratio of at least 3 were selected. The noise was calculated by the median absolute deviation. To eliminate mass drifts, the pipeline contains an alignment procedure using a kernel-based clustering approach adapted from pyBASIS^20^. Here peaks which were present in at least 3% of all pixels across all samples formed the common *m/z* vector to which the peaks were aligned by nearest-neighbour mapping. Then off-tissue/matrix pixels were identified and removed for each dataset (Figure S1). To identify matrix pixels, data of each sample were first reduced to two dimensions by using UMAP followed by HDBSCAN clustering. The cluster which was connected to most pixels of the border of the measured area was considered the matrix cluster. Clusters with a Spearman correlation above 0.7 to the matrix cluster were combined to an extended matrix cluster. In the third step the binary image of the extended matrix cluster was post-processed by removing isolated objects of up to 5 pixels followed by a binary closing operation using a 5×5 pixel square structuring element for the dilation and a 2×2 pixel square structuring element for the erosion. For the post-processing the scikit-image and SciPy libraries were used. After matrix removal the spatial coherence^43^ was calculated for each ion which measures it’s informativeness. Ions with low spatial coherence were removed. Then the data were normalised by median-fold-change (MFC) normalisation to account for intra-sample and inter-sample variation. To indicate spectra variations among the samples, a UMAP-based outlier detection method was applied (Figure S2). Here the data of all samples were first reduced to two dimensions by using UMAP followed by HDBSCAN clustering. Then sample-specific clusters (SSC) were identified. SSCs are clusters in which most pixels originate from one sample. Finally, samples in which most pixels were SSC pixels were considered sample outliers. De-isotoping was performed as last step. In an iterative approach, isotopes were identified based on their theoretical m/z value within a predefined tolerance range and their theoretical intensity pattern.

### Lipid annotation

In the first step, a list of tentative annotations of *m/z* values were made by using the bulk structure search provided on the LipidMaps website (www.lipidmaps.org). We searched for matches between all expected lipid classes (fatty acids/esters [FA], ceramides [Cer], sphingomyelins [SM], hexosyl ceramides [HexCer], triglycerols [TG], diglycerols [DG], glycerophosphocholines [PC], glycerophosphates [PA], glycerophosphoserines [PS], glycerophosphoethanolamines [PE], glycerophosphoglycerols [PG], sterols [ST]) and [M+H]^+^, [M+Na]^+^ and [M+K]^+^ precursor ions with a mass tolerance of +/- 0.01 *m/z*.

This list was then manually curated considering the biological context and expanded with the annotations gained from MALDI-DDA-MSI as well as other on-tissue MALDI MS/MS experiments. In addition, some lipids were validated by LC-MS/MS, as described in LC-MS/MS validation strategy for lipid annotations^64^. If no further information from MS/MS was gained, the annotation is only based on the lipid species level, which means in the context of ether lipids, that 1-O-alkyl lipids with at least one double bond could also be interpreted as 1-O-alkenyl ethers, as described in lipid nomenclature guidelines^65^.

### Data analysis, statistics and visualisation

Tissue segmentation was performed on the full m/z spectrum of all control and infected urinary bladder sections by UMAP (n_neighbors=10, min_dist=0.0, dist_metric=’cosine’) followed by HDBSCAN clustering (min_samples=50, min_cluster_size=10000). The clusters were manually merged into super-clusters which represent the main tissue context (urothelium, lamina propria and muscle).

Then mean spectra of the pixel clusters were extracted and statistically analysed using the Python package SciPy. The standard t-test was used for normally distributed populations with equal variances and the Welch’s t-test for normally distributed populations with unequal variances. Non-normally distributed populations were statistically analysed by the Wilcoxon rank-sum test. The Shapiro-Wilk test and Levene test was used to test for normal distribution and equal variances. The ratio between the means of two populations (here control and infected) was determined as log_2_(fold change).

For the lipidomic analysis of neutrophils, we reduced the *m/z* spectrum by filtering out *m/z*-values for which no lipid in the database could be matched based on our m/z tolerance, as well as tissue-specific *m/z*-values with a Pearson’s correlation (SciPy Python package) above 0.5 to one of the tissue regions extracted from the previous tissue segmentation (urothelium, lamina propria, muscle). To differentiate between Ly6G^+^ and Ly6G^-^ pixels, we segmented the registered Ly6G images using the Python package Scikit-image. First images were Gaussian smoothed with sigma=1 followed by Otsu thresholding. Finally small objects (below 10 pixels size) were removed. Unsupervised clustering of Ly6G^+^ pixels was performed by UMAP (n_neighbors=3, min_dist=0.0, dist_metric=’cosine’) followed by HDBSCAN clustering (min_samples=30, min_cluster_size=500).

Important lipids for the ROIs (here tissue regions and neutrophils) were extracted by using Extreme Gradient Boosting (XGBoost)-based classification with class weights, Pearson’s correlation and SHAP values. Therefore, the Python packages XGBoost, SciPy and SHAP were used. Class weights were calculated by #samples / (#classes * #occurences) using the Python package scikit-learn. The Python packages Pandas, Seaborn, Matplotlib and matplotlib-venn were used to generate all visualisations (volcano plots, scatterplots, pie charts, Venn diagrams, segmented images, and bar plots).

### Microscopy data pre-processing

Images were pre-processed by rolling-ball background subtraction, Gaussian smoothing with sigma=3 followed by percentile stretching using the Python library scikit-image.

### Registration of microscopy and MALDI MSI

Initial transformation of the images was performed in Fiji for microscopy (rotation and cropping) to match the imaged tissue region and orientation of both modalities. For precise registration we used symmetric normalisation (SyNRA) as transformation, consisting of rigid, affine and deformable transformation, and Mattes mutual information as optimisation metric implemented in the Advanced Normalisation Tools (ANTs) library^30^. We down sampled the microscopy image, using linear interpolation, to the same spatial dimensions as the MALDI image using the Python package ImgAug (https://github.com/aleju/imgaug). The down sampled AF from microscopy was used as moving image and the UMAP image from MSI, which visualises the main tissue structure similar to the AF image, was used as fixed image. We validated the registration result by the Jaccard index for the overlap between registered urothelial mask from microscopy and MSI mask. We applied spatial k-means clustering on the MSI data and selected the cluster containing the urothelium. The urothelial mask from microscopy was manually created.

## Data availability

The intermediate pre-processing results of the MALDI-2 MSI bladder data are available upon request due to data size. All other (t-)MALDI-2 MSI and IFM data (raw and generated results) were deposited at Zenodo (https://doi.org/10.5281/zenodo.11913042).

## Code availability

The msiFlow source code is publicly available on GitHub (https://github.com/Immunodynamics-Engel-Lab/msiflow).

## Ethics statement

The animal study was approved by LANUV, Recklinghausen, Germany. The study was conducted in accordance with the local legislation and institutional requirements.

## Author contributions

P.S. designed the software. P.S. developed and implemented the algorithms and workflows. D.S. implemented the browser-based user interface. L.M. implemented the raw MSI data reader. S.B. acquired MSI data. L.W., J.B., M.R. and S.T. acquired IFM data. L.W., J.B. and S.T. performed animal experiments and sample preparation. M.R. performed isolation of neutrophils. S.S.K. performed LC–MS/MS measurements. P.S., S.B., L.W., J.S., O.Sh., K.D., P.P., S.S.K. and D.R.E. interpreted the data. S.D.K, H.H., D.F., R.V.P., L.C.M., M.G., O.So. contributed to the project development. P.S. and D.R.E. wrote the manuscript. D.R.E. supervised and coordinated the study. All authors revised the manuscript and approved the submitted version.

## Acknowledgments

We acknowledge support by the Open Access Publication Fund of the University of Duisburg-Essen, the Central Animal Facilities of the Medical Faculty Essen, the Imaging Center Essen (Alexandra Brenzel and Dr. Anthony Squire) and the Immunoproteomics group (Dr. Olga Shevchuk, Stephanie Tautges-Schaefer, Stephanie Thiebes and Jenny Dick), supported by INST 20876/486-1 to DRE.

