## Supplementary for "msiFlow: Automated Workflows for Reproducible and Scalable Multimodal Mass Spectrometry Imaging and Immunofluorescence Microscopy Data Processing and Analysis"

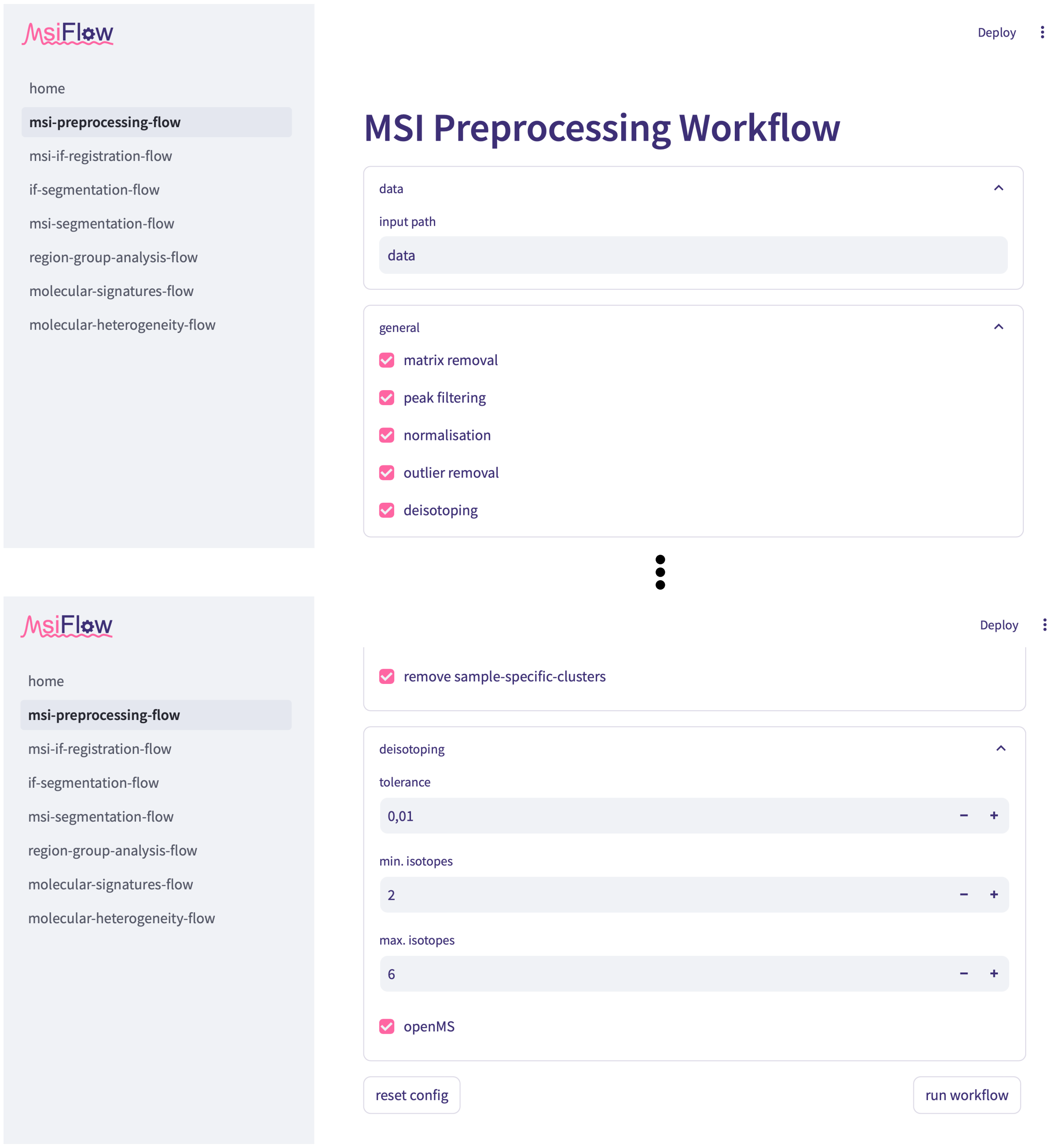


**Supplementary Fig. 1: Representative page of one of the workflows of the browser-based user interface of msiFlow.** In the browser-based user interface of msiFlow a workflow can be selected from the sidebar on the left. Here the pre-processing workflow is selected. The workflow page allows to edit the workflow parameters. The lower left button resets the configuration parameters to the default options. The lower right button executes the workflow. The browser-based version of msiFlow is available on GitHub (https://github.com/Immunodynamics-Engel-Lab/msiflow).

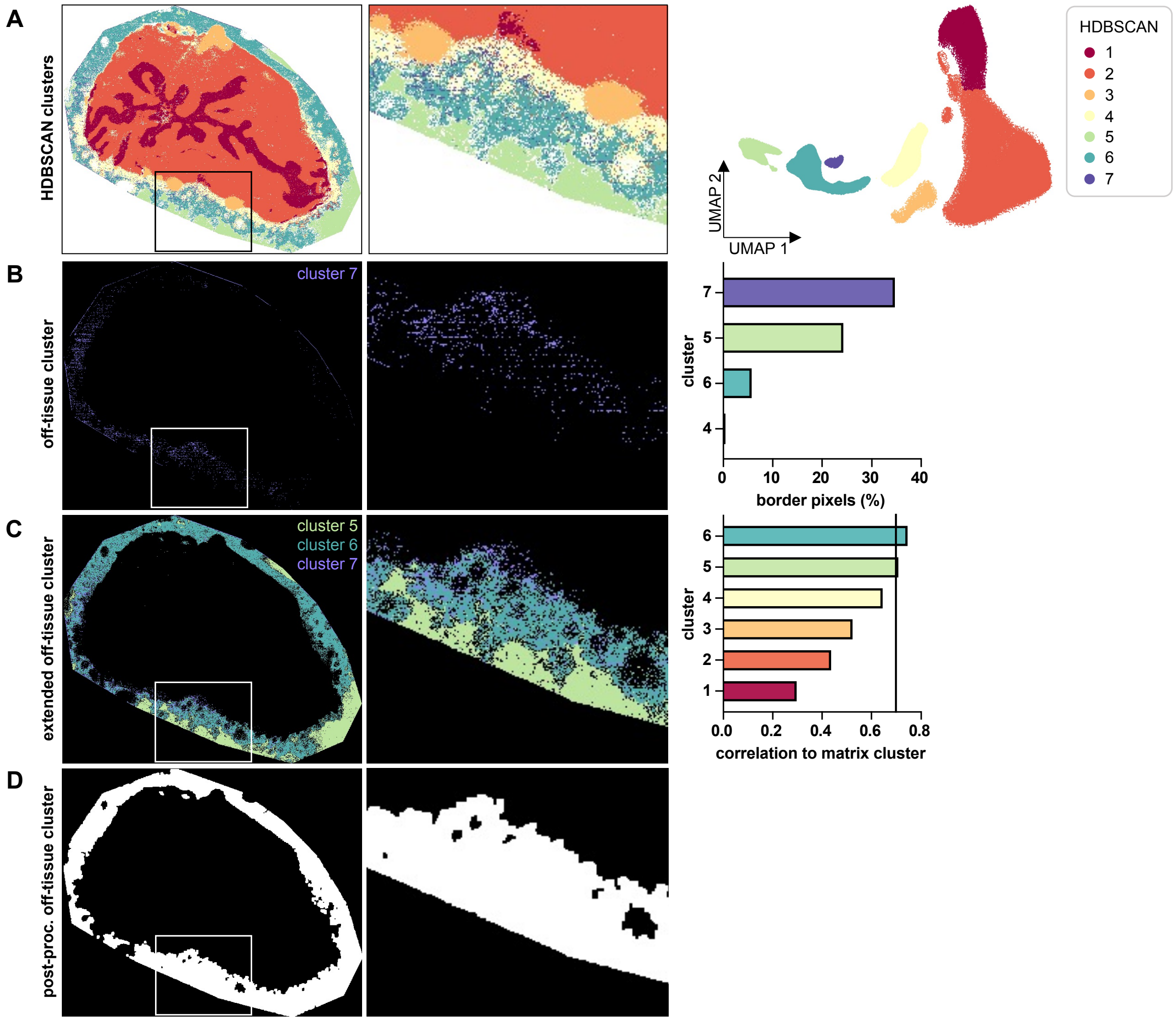


**Supplementary Fig. 2: Identification of off-tissue pixels. Related to Figure 1B5. (A)** Data were reduced to 2 dimensions by UMAP followed by HDBSCAN clustering. **(B)** The cluster which was connected to most border points of the measured area (here cluster 7) was considered off-tissue cluster. **(C)** Clusters with a Spearman correlation above 0.7 to the off-tissue cluster (indicated by the vertical line in the bar plot) were combined to an extended off-tissue cluster. **(D)** Post-processing of the extended off-tissue cluster (Fig. S2C) by removing objects of up to 5 pixels followed by a binary closing operation using a 5x5 pixel square structuring element for the dilation and a 2x2 pixel square structuring element for the erosion.

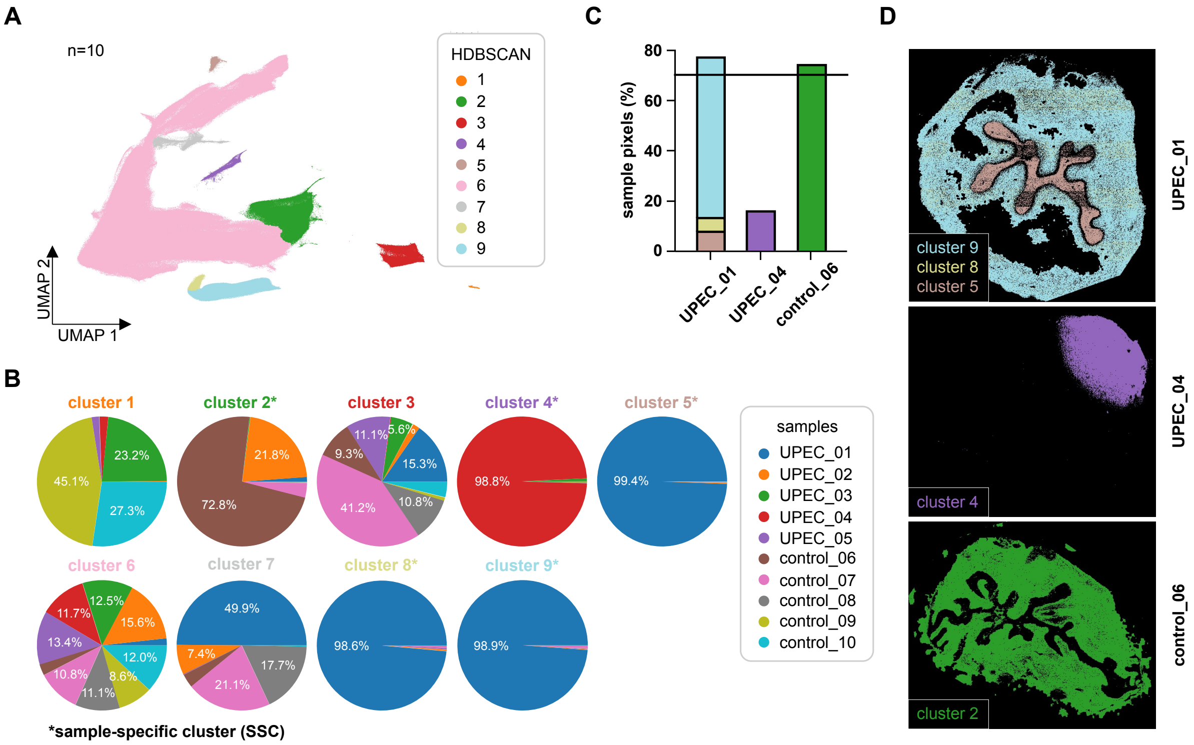


**Supplementary Fig. 3: Clustering algorithm to identify spectra variations among the samples. Related to Figure 1B8. (A)** Data of all samples were reduced to 2 dimensions by UMAP to indicate the main clusters identified by HDBSCAN clustering. **(B)** Pie charts indicating samples from each cluster in percentages. Clusters in which most pixels (here at least 70%) originated from one sample were considered sample-specific clusters (SSC). **(C)** Bar plot showing the percentage of pixels covered by SSC pixels for each sample containing SSCs. The sample was considered an outlier if 70% of the pixels in a sample were SSC pixels. **(D)** Spatially resolved visualization of the SSCs.

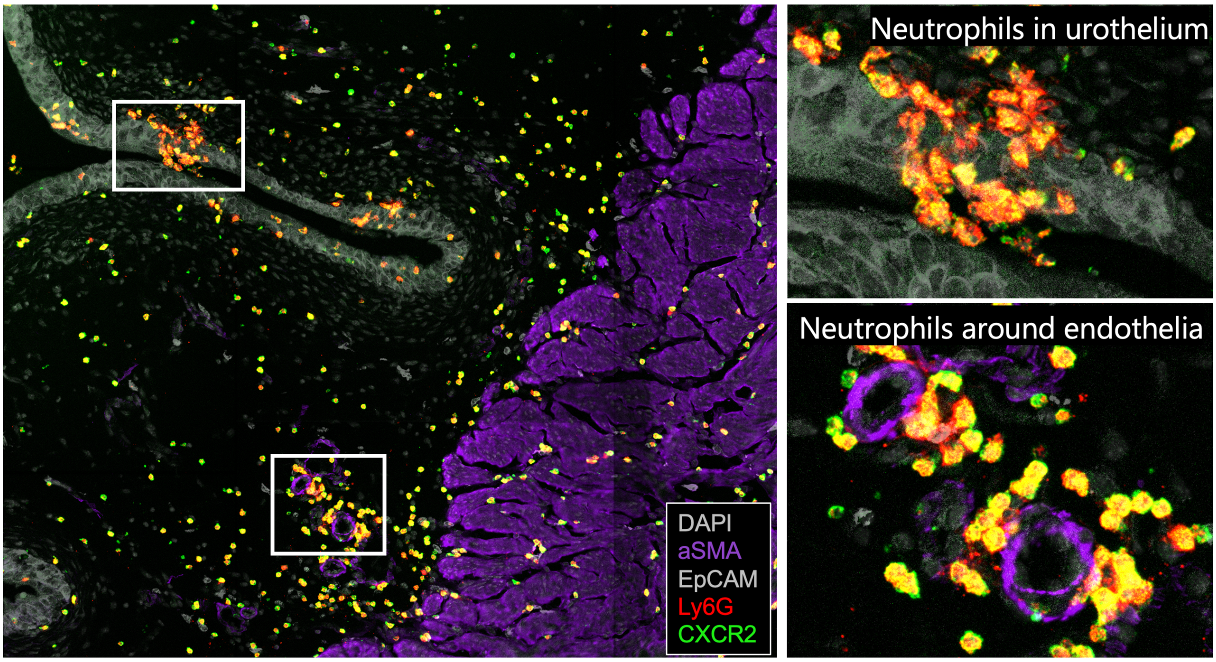


**Supplementary Fig. 4: Spatial regulation of CXCR2 by neutrophils in urinary tract infection.** MACSima image of urinary bladders one day after infections showing neutrophils (Ly6G) around blood vessels (aSMA; lower quadrant in image to the left and lower image to the right) in the lamina propria, and in the urothelium (EpCAM, top image to the right).

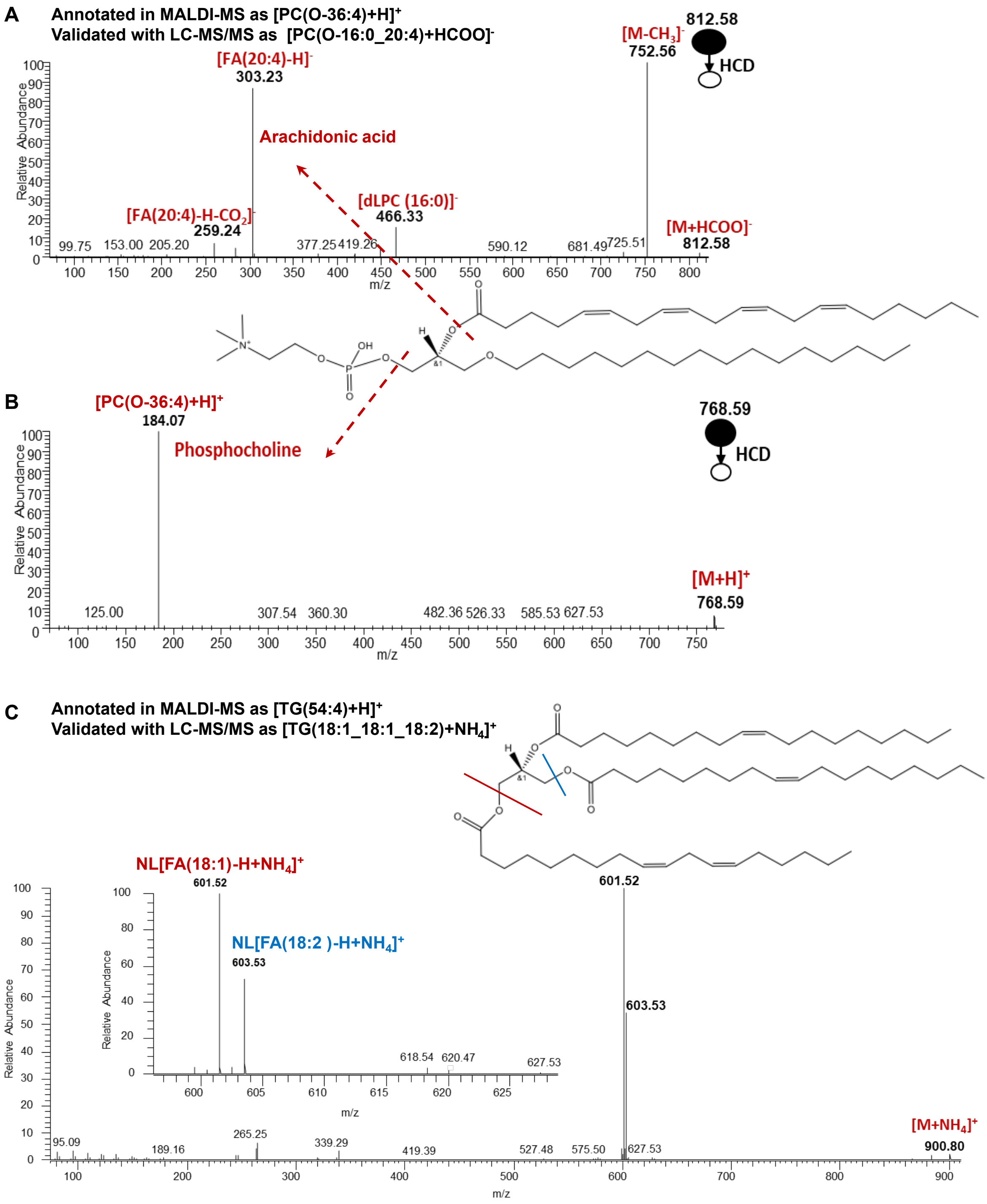


**Supplementary Fig. 5: Validation of lipid annotations by liquid chromatography-tandem mass spectrometry (LC-MS/MS) in negative- and positive-ion mode.** High-energy collisional dissociation fragmentation (HCD-MS/MS) spectra of **(A)** [PC(O-36:4)+HCOO]$^-$ *m/z* 812.5811 and **(B)** [PC(O-36:4)+H]$^+$ *m/z* 768.5902, NL; neutral loss, FA; fatty acid, dLPC; demethylated lysophosphatidylcholine. **(C)** Annotation with MALDI MSI as [TG(54:4)+H]$^+$ was validated as [TG(18:1_18:1_18:2)+$NH_4$]$^+$ with LC-MS/MS.

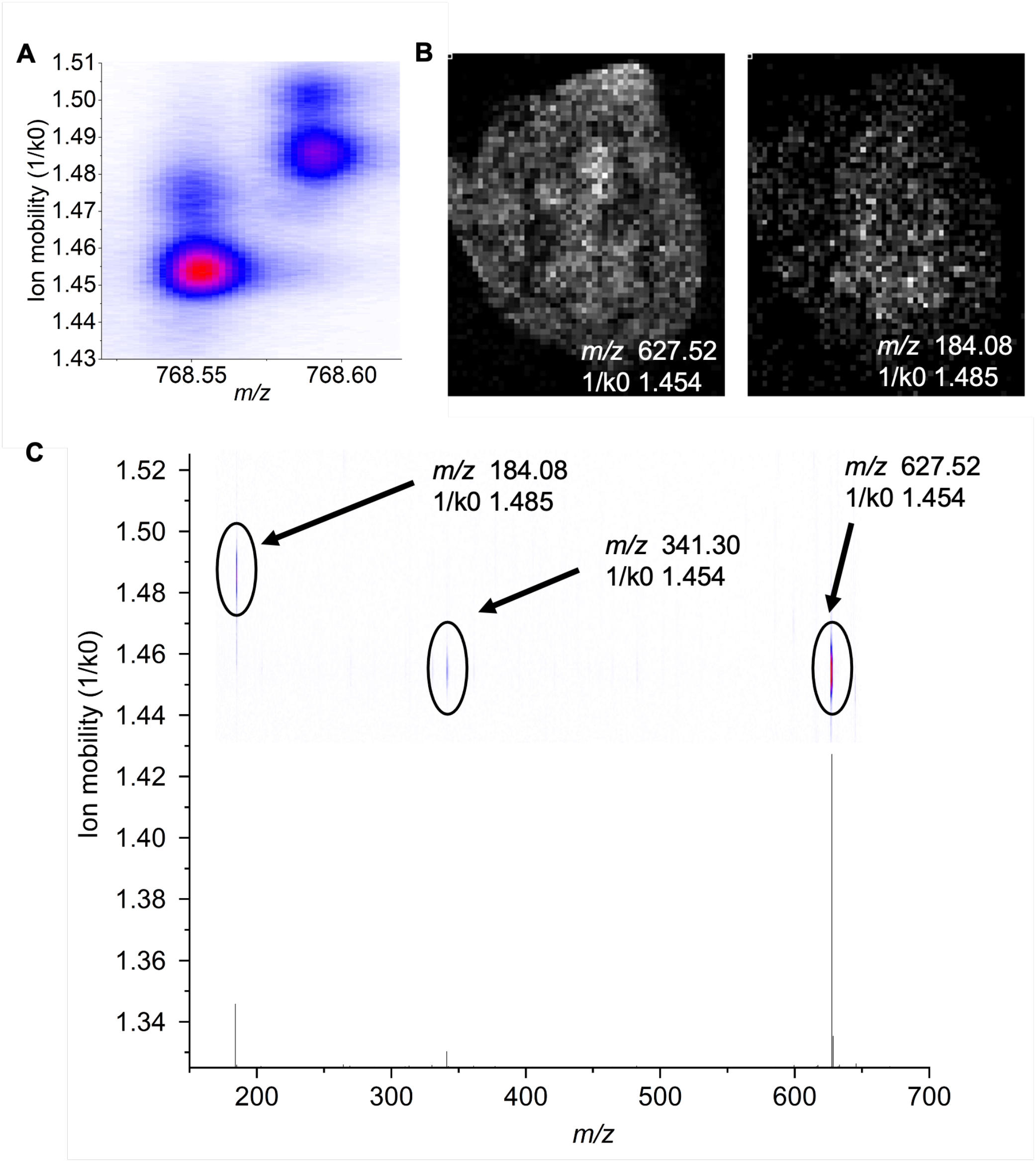


**Supplementary Fig. 6: On-tissue MS/MS measurement of 768.6 *m/z* precursor ion in Tims ON mode. (A)** Ion mobility separation of 768.55 m/z and 768.60 m/z. **(B)** Ion images of 627.52 *m/z* (left) and 184.08 *m/z* (right). **(C)** Ion mobility of fragment ions**.**

A

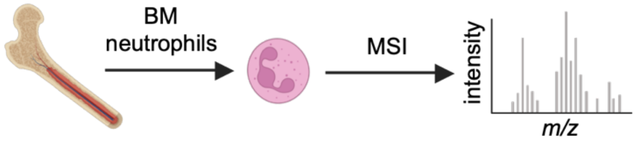


B

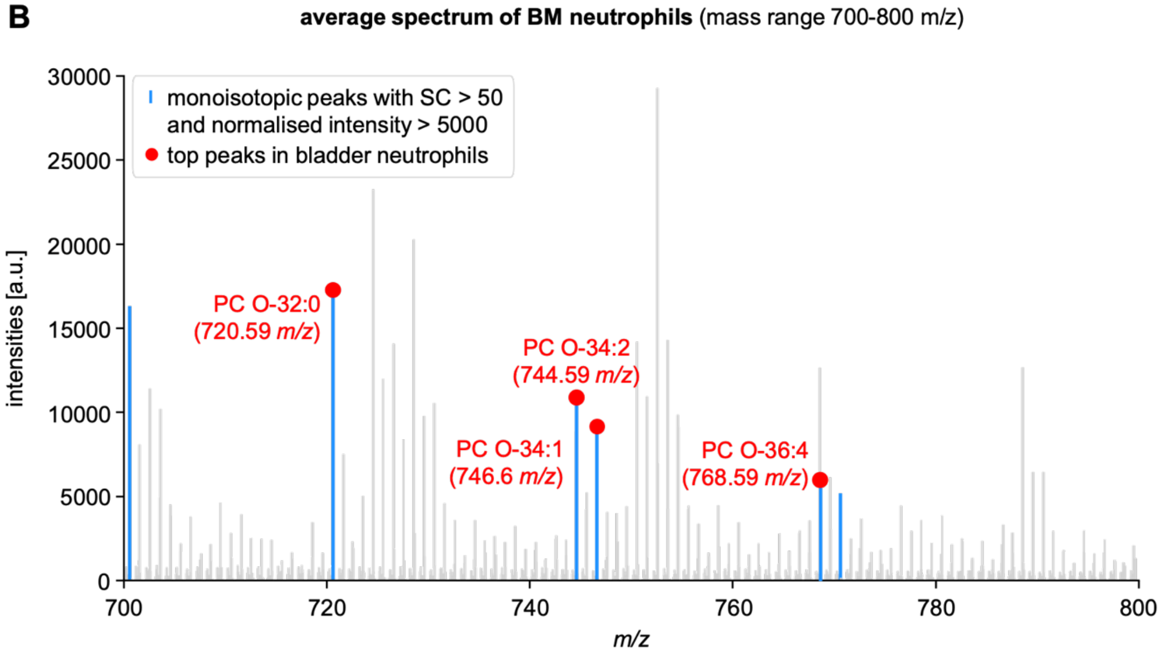


**Supplementary Fig. 7: Validation of identified peaks for neutrophils by MSI of bone marrow neutrophils. (A)** Neutrophils were isolated from murine bone marrow followed by MSI analysis. **(B)** Average spectrum of BM neutrophils highlighting the monoisotopic peaks with a spatial coherence of at least 50 and normalised intensity of at least 5000 (blue). The top 4 lipids (+H) for bladder neutrophils are highlighted in red. BM (bone marrow), spatial coherence (SC).

**Supplementary Table 1: Parameters used by msiFlow for pre-processing MALDI MSI data.**

| Parameter | Type | Value | Description |
|---|---|---|---|
| ***smoothing and peak picking*** | | | |
| **snr** | numeric | 3 | signal-to-noise threshold for peaks |
| **smooth** | numeric | 1 | set to 1 to perform Savitzky-Golay smoothing |
| **window** | numeric | 11 | length of the Savitzky-Golay filter window |
| **order** | numeric | 3 | order of the polynomial of Savitzky-Golay filter |
| ***alignment*** | | | |
| **num_pixel_percentage** | percentage | 80 | percentage of pixels to consider for building common m/z vector |
| **mz_resolution** | numeric | 0.005 | bin size in Da of histogram which is used to build common m/z vector |
| **pixel_percentage** | percentage | 3 | min. percentage of m/z to form common m/z vector |
| **max_shift** | numeric | 0.01 | max. shift in Da to shift peaks |
| ***matrix removal*** | | | |
| **clustering** | boolean | True | set to True to use clustering for matrix/off-tissue identification |
| **dim_reduction** | categorical | umap | method to reduce spectra, either umap, t-sne or pca |
| **n_components** | integer | 2 | number of components of dim. reduction |
| **metric** | numeric | cosine | distance metric of UMAP |
| **n_neighbors** | integer | 100 | size of local neighborhood UMAP will look at |
| **min_dist** | numeric | 0.0 | min. distance apart that points are allowed to be in low dim. representation (UMAP) |
| **cluster_algorithm** | categorical | hdbscan | algorithm to cluster low dim. representation, either hdbscan, k-means or gaussian_mixture |
| **min_cluster_size** | integer | 1000 | min. size of an HDBSCAN cluster |
| **min_samples** | integer | 500 | min. number of neighbors to a core point (HDBSCAN) |
| **matrix_corr_thr** | numeric | 0.7 | clusters with this Spearman correlation threshold to the initial off-tissue cluster are combined to an extended matrix cluster |
| **pixel_perc_thr** | numeric | 30 | pixel percentage threshold of clusters to extend off-tissue cluster |
| **matrix_postproc** | boolean | True | set to True for post-processing of the matrix/off-tissue image |
| **pixel_removal** | boolean | True | set to True to remove matrix/off-tissue pixels from each dataset |
| **matrix_subtraction** | boolean | False | set to True to subtract the mean matrix spectrum from each pixel spectrum |

| | | | |
|---|---|---|---|
| **num_matrix_peaks** | numeric | 0 | set number of top matrix peaks to remove |
| **matrix_peak_removal** | boolean | False | set to True to remove matrix peaks |
| **matrix_mzs** | list | “” | list of known matrix m/z values |
| | | ***intra-normalisation*** | |
| **method** | categorical | mfc | method for intra-sample normalisation: either sum, mean, median or mfc |
| | | ***inter-normalisation*** | |
| **method** | categorical | mfc | method for inter-sample normalisation: either sum, mean, median or mfc |
| | | ***peak filtering*** | |
| **sum** | categorical | max | method to summarize the spatial coherence over all samples, either min, mean or max |
| **thr** | numeric | 1000 | filter peaks based on this defined spatial coherence threshold |
| | | ***sample outlier detection*** | |
| **n_neighbors** | integer | 10 | size of local neighborhood UMAP will look at |
| **min_cluster_size** | integer | 5000 | min. size of an HDBSCAN cluster |
| **min_samples** | integer | 1000 | min. number of neighbors to a core point (HDBSCAN) |
| **cluster_thr** | percentage | 70 | cluster pixel percentage which must be covered by one sample to be considered a sample-specific cluster (SSC) |
| **sample_thr** | percentage | 70 | sample pixel percentage which must be covered by SSC pixels to be considered a sample outlier |
| **remove_ssc** | boolean | True | set to True to remove SSC pixels |
| | | ***de-isotoping*** | |
| **tolerance** | numeric | 0.01 | the tolerance used to match isotopic peaks |
| **min_isotopes** | integer | 2 | the min. number of isotopic peaks |
| **max_isotopes** | integer | 6 | the max. number of isotopic peaks |
| **openMS** | boolean | False | If True, openMS routine is applied |

**Supplementary Table 2: Lipid identification by DDA**

| m/z | Name | ppm |
|---|---|---|
| 569.5152 | DG(32:0) | 2.1 |
| 690.5088 | PE(32:1) | 2.9 |
| 703.5759 | SM(34:1) | 1.5 |
| 704.5248 | PE(33:1) | 3.3 |
| 718.5395 | PE(34:1) | 1.8 |
| 720.5906 | PC(O-32:0) | 0.6 |
| 730.5752 | PE(O-36:2) | 0.9 |
| 731.6075 | SM(36:1) | 1.8 |
| 732.5552 | PC(32:1) | 1.9 |
| 734.5705 | PC(32:0) | 1.5 |
| 740.5239 | PE(16:0_20:4) | 1.9 |
| 744.5552 | PE(36:2) | 1.9 |
| 744.5900 | PC(O-34:2) | 0.2 |
| 746.5707 | PE(36:1) | 1.6 |
| 746.6058 | PC(O-34:1) | 0.0 |
| 748.5289 | PE(O-38:7) | 1.7 |
| 758.5706 | PC(34:2) | 1.5 |
| 759.6388 | SM(38:1) | 1.8 |
| 760.5863 | PC(34:1) | 1.6 |
| 762.5301 | PS(34:1) | 2.8 |
| 762.5999 | PC(34:0) | 1.1 |
| 764.5237 | PE(38:6) | 1.5 |
| 766.5393 | PE(38:5) | 1.5 |
| 768.5551 | PE(18:0_20:4) | 1.7 |
| 782.5701 | PC(36:4) | 0.8 |
| 784.5846 | PC(36:3) | 0.6 |
| 786.6017 | PC(36:2) | 1.3 |
| 787.6699 | SM(40:1) | 1.5 |
| 788.5452 | PS(36:2) | 2.0 |
| 790.5617 | PS(36:1) | 3.0 |
| 792.5548 | PE(40:6) | 1.3 |
| 796.5861 | PE(40:4) | 1.3 |
| 810.6022 | PC(38:4) | 1.8 |
| 811.6709 | SM(42:3) | 2.7 |
| 812.5450 | PS(38:4) | 1.7 |
| 813.6855 | SM(42:2) | 1.4 |
| 815.6997 | SM(42:1) | 0.4 |
| 836.5451 | PS(40:6) | 1.8 |
